# Classification of containment hierarchy for point clouds in periodic space

**DOI:** 10.1101/2025.08.06.668936

**Authors:** Bart M. H. Bruininks, Ilpo Vattulainen

**Author notes:** BMHB designed the research, developed the mathematics, performed the programming and wrote the manuscript. ITV helped with the manuscript revisions. Contributing authors.

## Abstract

When working with large point clouds, it is often useful to label the data. One of such labels is the classification of insides and outsides. Meaning that as input an a selection can be given and the output will be all that lies inside that selection. Containment labeling has shown to be very useful for point clouds in Euclidean space and allows for the generation of signed distance fields, however, such classification was not robustly available in periodic space.

Here were present our open source tool coined MDVContainment, which rigorously solves the containment problem for the periodic case. This algorithm was applied to a coarse grained acyl chains bicelle, transfecting lipoplex and a system of stacked bilayers as well as to an all atomistic periodic nanotube. Showing that the analysis is performing well—both atomistic and coarse grained systems. The containment processing of these systems takes roughly the same amount of time as creating their universe objects in MDAnalysis (i.e. reading in the data).

Having a rigorous definition of containment makes it possible for MD analysis/visualization tools to support periodic containment, just as it is supported for non-periodic spaces.

## 1 Introduction

Point clouds are a common data structure to deal with volumetric data in a sparse manner. In molecular dynamics (MD) point clouds are used to represent particles in the simulation. As the computational capability increases, so are the sizes of the systems and their corresponding point clouds. Nowadays point clouds with millions of data points per frame are becoming more and more common[1–4]. In all atomistic (AA) MD this corresponds to a small lipid vesicle in solution (∼20 nm in diameter), for Martini CG simulations a 10M bead system is in the range of a experimental size vesicle (∼60 nm in diameter). Such systems are large enough to display complex nested behavior of compartmentalization and the notion of inside and outside becomes very useful.

It is rather intuitive for humans to talk about containment, we often assume the content to come with the container, unless explicitly stated otherwise. To give an example, when someone asks you to hand over the inflated balloon, we usually intend to ask for the balloon plus all the air it contains. No need to empty the balloon of air before handing it over. Unsurprisingly, the notion of containment has been studied extensively and for non-periodic regular spaces the problem has a nice solution (Jordan–Brouwer theorem[5]). This states that any (*n*−1)-sphere in n dimensions (*n* > 0) splits the space into two connected components, a bound component (the inside) and an unbound component (the outside). However, in periodic space (often used in MD) not all surfaces which separate two components are an n-sphere at all, nor does slicing with this boundary need to result in a bounded and unbounded connected component. This complicates the matter of containment, which is addressed and solved in this manuscript. The final containment algorithm was implemented using Python and Cython and was validated on both constructed and real MD data.

## 2 Theory

We will start by clarifying the objective and what we mean by containment. To do so we will explicitly state the goal and eight axioms. We finalize the theory section with the containment algorithm pseudo-code.

The goal is the classifier we wish to obtain. The axioms are an extension of well-known mathematical concepts. Concepts such as connected components[6] and percolation[7] are taken for granted. Each axiom comes with a figure, to enforce what is intended.

### 2.1 Definitions

#### Goal

To obtain a definition of containment resulting in a containment graph spanning the complete space. The containment graph is a directed acyclic (multi) graph. The components are the nodes in the graph and the directions of the edges indicate containment. The graph should satisfy the following conditions: First, a node cannot have more than one inward pointing edge. A, node can have more than one outward pointing edge. Second, the alteration of the containment hierarchy downstream should never affect the containment hierarchy upstream. All these conditions are satisfied by the axioms below and result in unambiguous specification of the containment hierarchy. An example of such a containment graph is given in Fig 1.

**Fig. 1.**
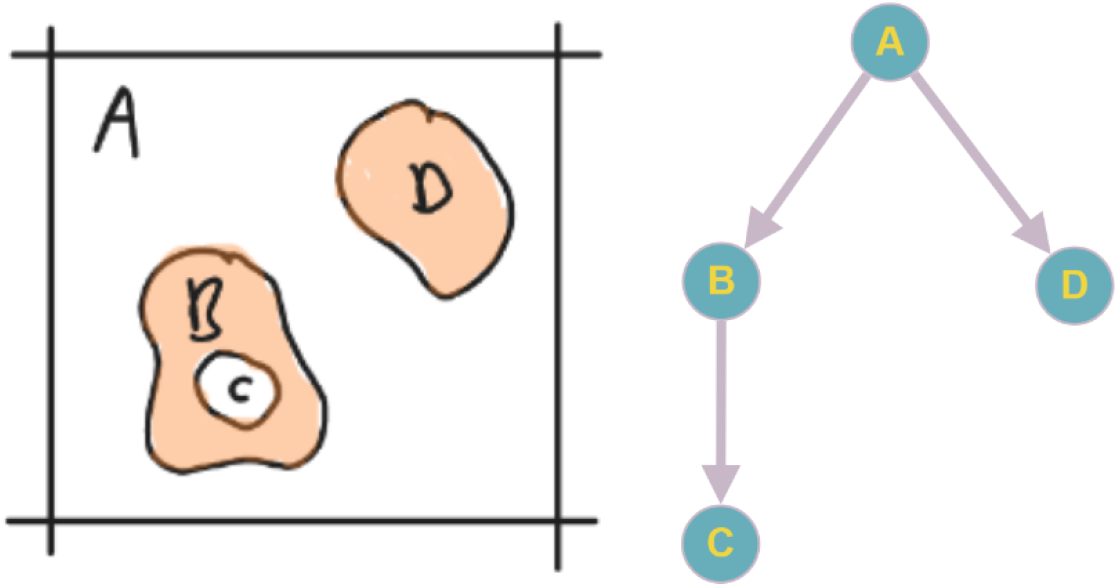
A containment graph of four components in 2D periodic space (ℝ^2^/ℤ^2^). A component contains all downstream components in the containment graph, plus itself. A single component (*A*) contains all the other components ({*A, B, C, D*}). There is one nested container (*B*) which also contains a component (*C*). The graph was drawn using https://graphonline.ru/en/.

#### Axiom 1. space

Space is regular and not curved (e.g. ℝ, ℤ, ℝ/ℤ, etc.; Fig. 2).

**Fig. 2.**
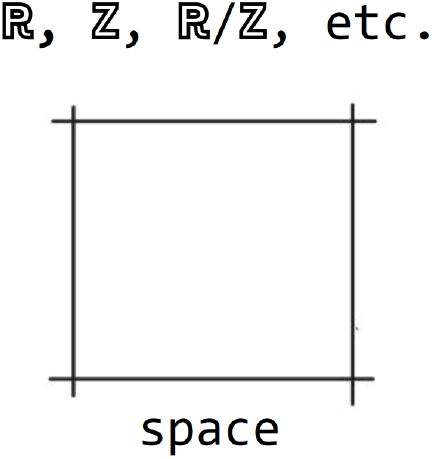
Axiom 1. *space*. The space is drawn as square with a boundary. It needs not be a square nor does it has to have a boundary, this is just for illustrative purposes.

#### Axiom 2. components

The binary partitioning of space results in connected components which can be *True* or *False*. Whenever we use the word *component* we mean *connected component* unless stated otherwise (Fig. 3).

**Fig. 3.**
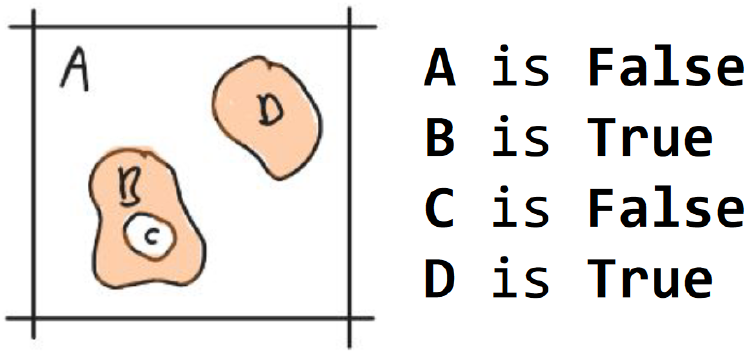
Axiom 2. *components*. The components B, C and D are contained, A is not. The densities{*A, D*} are represented in peach. The void components {*A, C*} are white.

#### Axiom 3. complements

The complement of a component is all of space which is not the component, therefore a complement does not need to be connected. The complement of a component *A* is annotated as !*A* (Fig. 4).

**Fig. 4.**
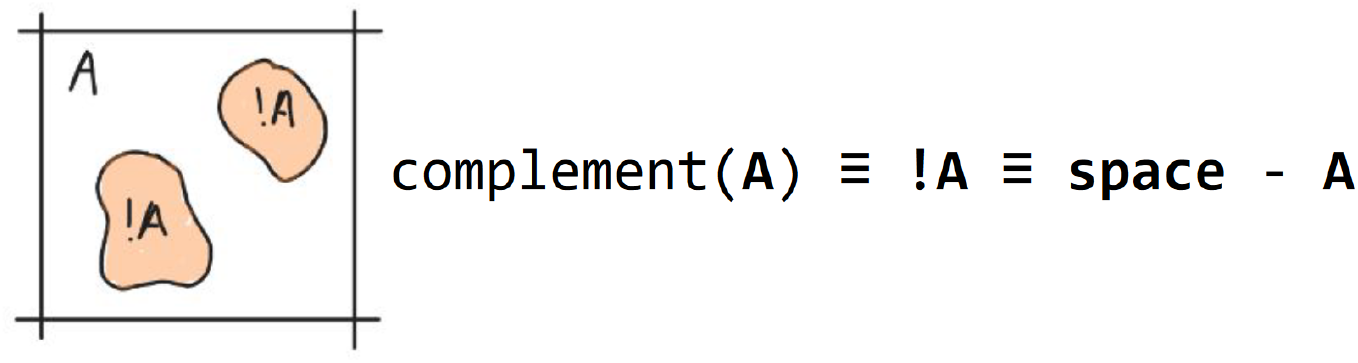
Axiom 3. *complements*. The complement !*A* is drawn in peach and its corresponding component *A* is drawn in white.

#### Axiom 4. rank

The rank (*ρ*) of a component, or its complement, is defined by the amount of orthogonal dimensions in which it percolates space [7]. The rank of a component or complement (*A* of rank 2) is indicated with a preceding subscript number (_2_*A*). Since a complement does not need to be connected, the rank of a compliment is the highest rank among the ranks of its components (Fig. 5).

**Fig. 5.**
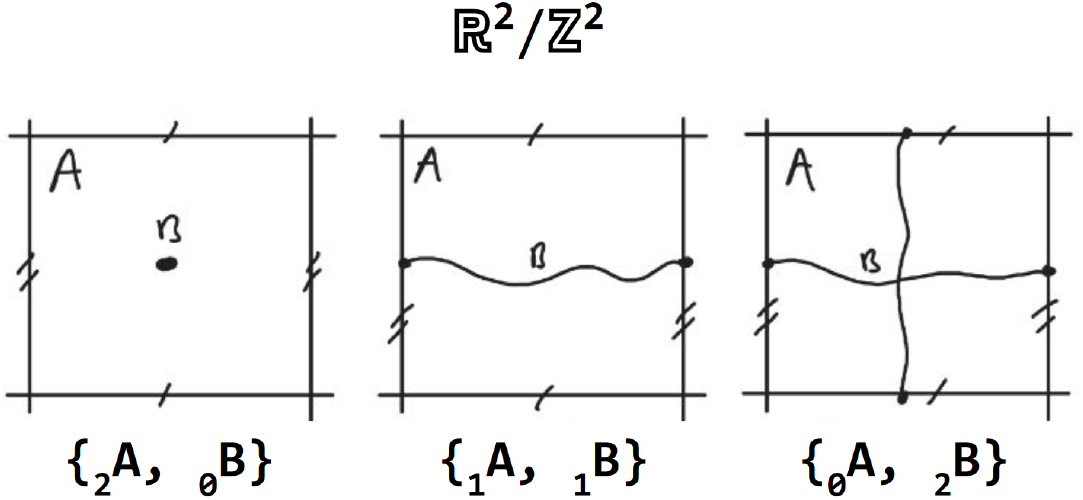
Axiom 4. *rank*. The rank of the void *A* with a dot (2), periodic line (1) and periodic net (0) are shown from left to right. The corresponding density *B* is of rank 0, 1 and 2 reciprocally.

#### Axiom 5. component containment

A component is contained if its complement is of higher rank. The containment status of a component (*A*) is annotated with a subscript *c* (*A*_*c*_; Fig. 6).

**Fig. 6.**
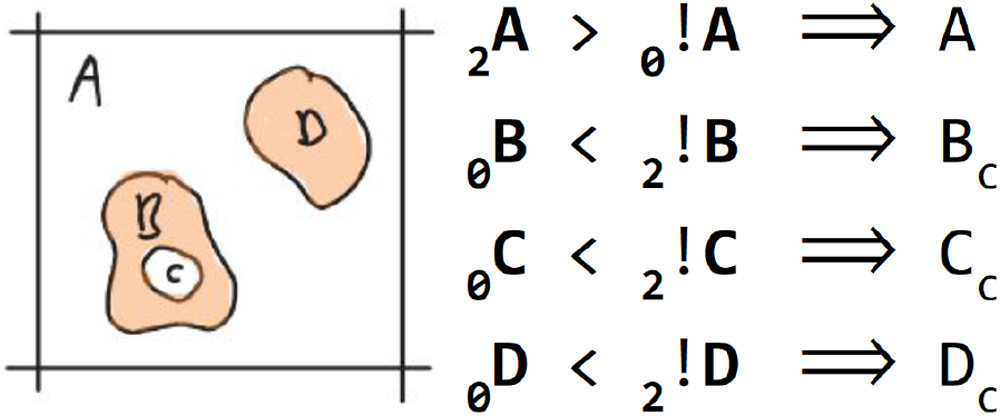
Axiom 5. *component containment*. The components {*B, C, D*} are contained, *A* is not. The densities {*A, D*} are represented in peach. The void components {*A, C*} are white).

#### Axiom 6. component contacts

Two components are in contact if they share a boundary. A contact between a component (*A*) and another component (*B*) is annotated with a tilde (*A*∼*B*). The collection of all contacts is called a contact graph (Fig. 7).

**Fig. 7.**
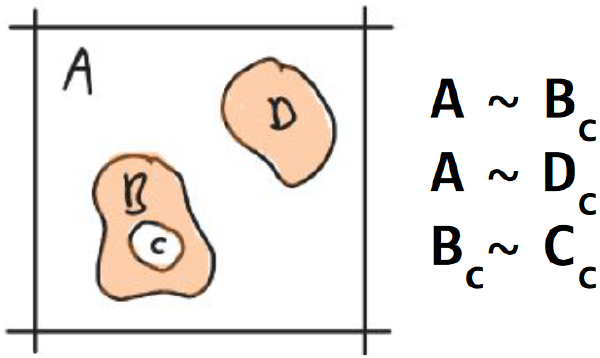
Axiom 6. *component contacts*. Component *A* is in contact with components {*B, D*} and *B* is in contact with *C*. The densities {*A, D*} are represented in peach. The void components {*A, C*} are white. The boundaries are represented with black curves.

#### Axiom 7. neighbor containment

A component is contained by a neighboring component in the contact graph if it is contained and the neighbor is not. Containment of a component (*A*) by another component (*B*) is annotated with an arrow (*B*→*A*). The collection of all containment hierarchy in a space is called its containment graph (Fig. 8).

**Fig. 8.**
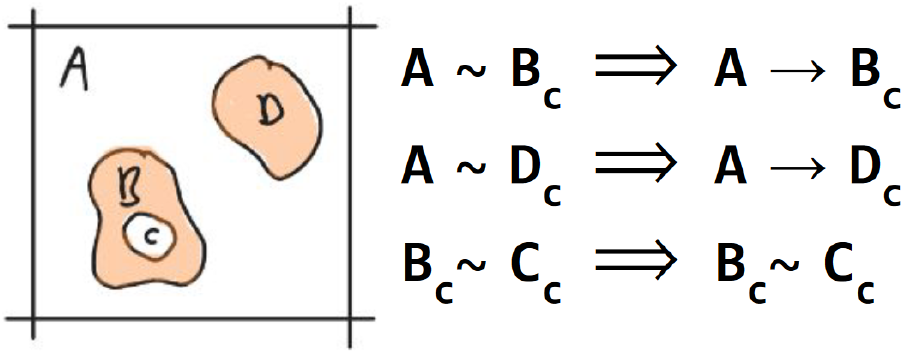
Axiom 7. *neighbor containment*. Component *A* contains *B* and *D*. The densities {*A, D*} are represented in peach. The void components {*A, C*} are white.

#### Axiom 8. recursive neighbor containment

A component which is contained by a neighboring component contains all other components it is in contact with (Fig. 9).

**Fig. 9.**
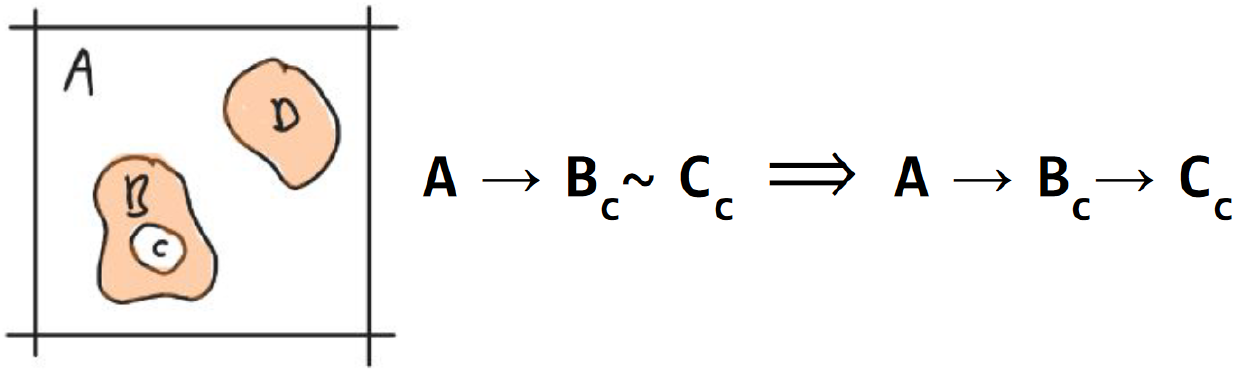
Axiom 8. *recursive neighbor containment*. Component *C* is contained by *B* due to recursion. The densities {*A, D*} are represented in peach. The void components {*A, C*} are white.

### 2.2 Pseudo code

#### Algorithm 1 Containment Analysis Algorithm

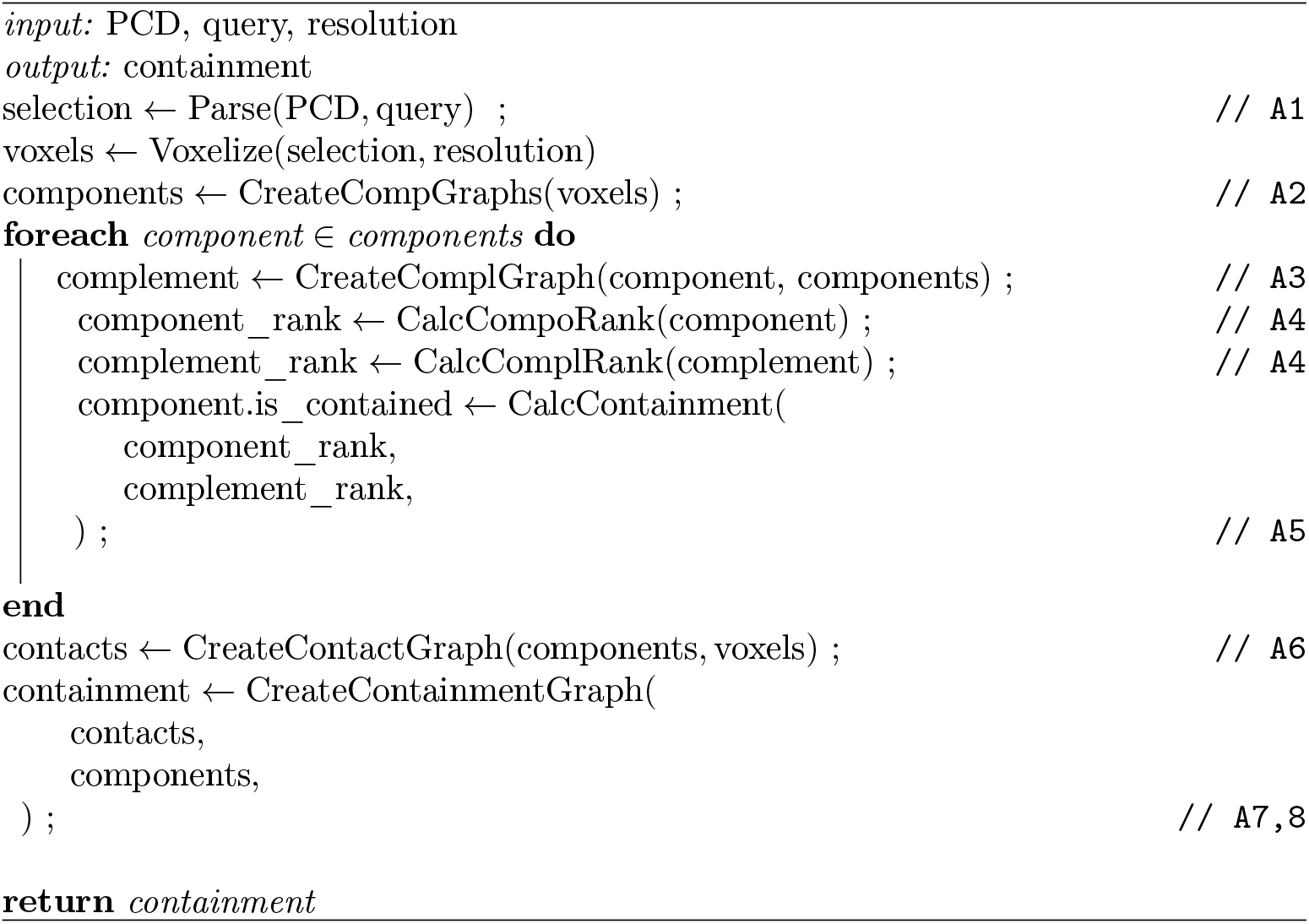

## 3 Results

The containment algorithm itself is the center of this work and the applications to the variety of systems are there to make the reader appreciate its scope. We start our results with synthetic 2D examples to enforce the intuition for what is meant by containment. Finally, we show containment results for real 3D MD systems.

### 3.1 Containment in ℝ^2^*/*ℤ^2^

The periodic space is often used in molecular dynamics to alleviate boundary effects. We will therefore show some examples in such a space using a discretized framework of connectivity. The discretized volumes are pixels and each pixel is connected to its eight neighbors with which it shares a vertex. A pixel is set to *True* if a point lies inside its boundaries and it is set to *False* otherwise. We will call *True* components *densities* and *False* components *voids*.

To illustrate the proposed containment algorithm on such systems, we give five examples (Fig. 10). All graphs shown are created using *MDVContainment, VMD, Pyvis* and *Inkscape*.

**Fig. 10.**
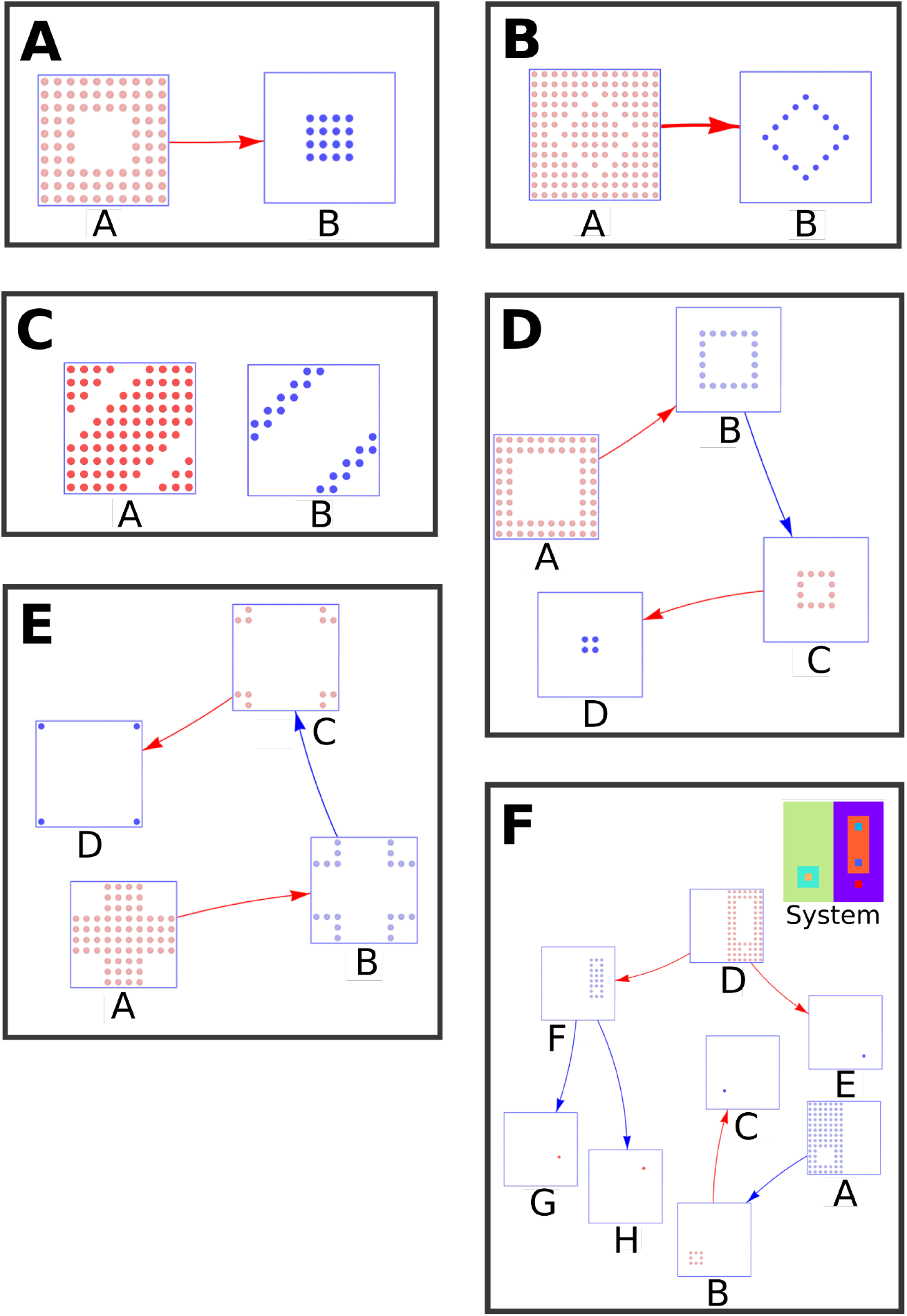
Containment hierarchy for ℝ^2^*/*ℤ^2^ synthetic data. (A) Single island. (B) Unintuitive containment due to bleeding over diagonal connectivity. (C) Diagonal coexistence of two rank 1 components.(D) Nested islands. (E) Translated nested islands. (F) A complex example of coexistent nested containers with branching; its components are shown in the top right. In the containment graphs, densities are blue and voids are red.

The first example (Fig. 10 A) shows a single density component (_0_*A*) in a void (_2_*B*). The complement of the density (_2_!*A*) is of higher rank than the density, so it is contained (*A*_*c*_). The complement of the void (_0_!*B*) is of lower rank than the void itself, therefore the void is not contained (*B*). The contained density is in contact with the non-contained void (*A*_*c*_∼*B*), thus the density is contained by it (*B*→*A*).

The second example (Fig. 10 B) contains a hollow diamond shaped density (_0_*A*) with a single void (_2_*B*) on both sides – due to diagonal connectivity. This results in the same contact graph and containment as shown in the first example (*B*→*A*). The density is too thin with respect to the chosen discretization resolution and unexpected, and probably unwanted, bleeding can occur. The lesson learned is that in a disrcretized space one should be careful when details are as small as the discretization volume. ‘Gaussian Blurring’ or ‘Binary Dilation/Closing’ of the density can alleviate this issue.

Third (Fig. 10 C), a diagonal rod density (_1_*A*) embedded with a void of (_1_*A*). Since both components have a complement (_1_!*A*, _1_!*B*) of equal rank neither of them is contained thus, no containment hierarchy is present at all.

Forth and fifth (Fig. 10 D, E), nested containers behave as expected and translation of the system does not alter the containment (up to discretization). The same holds for rotation (not illustrated). This is a direct consequence of the invariance of the definition of containment towards these properties.

The final example is a complex system in which multiple outsides exist, each with their respective containment hierarchy. This yields two disconnected sub-graphs in the containment hierarchy. Meaning, locally there might be a containment hierarchy even though the total space is fragmented.

### 3.2 Containment in ℝ^3^*/*ℤ^3^ MD data

As for the 2D case we will discretize our 3D space, this time resulting in voxels. A voxel is connected to its 26 neighbors with which it shares a vertex. A voxel is set to *True* if a point lies inside its boundaries and it is set to *False* otherwise. The lipid tail particles are used as the input point cloud for the discretization and we use a discretization size of 0.5 nm, performing one round of binary closing after discretization. This to close small pores in continuous density, caused by the discretization, due to the sparseness of the Martini coarse grained particles. For the atomistic data (Fig. 11 B) no closing was needed at a resolution of 0.5 nm.

**Fig. 11.**
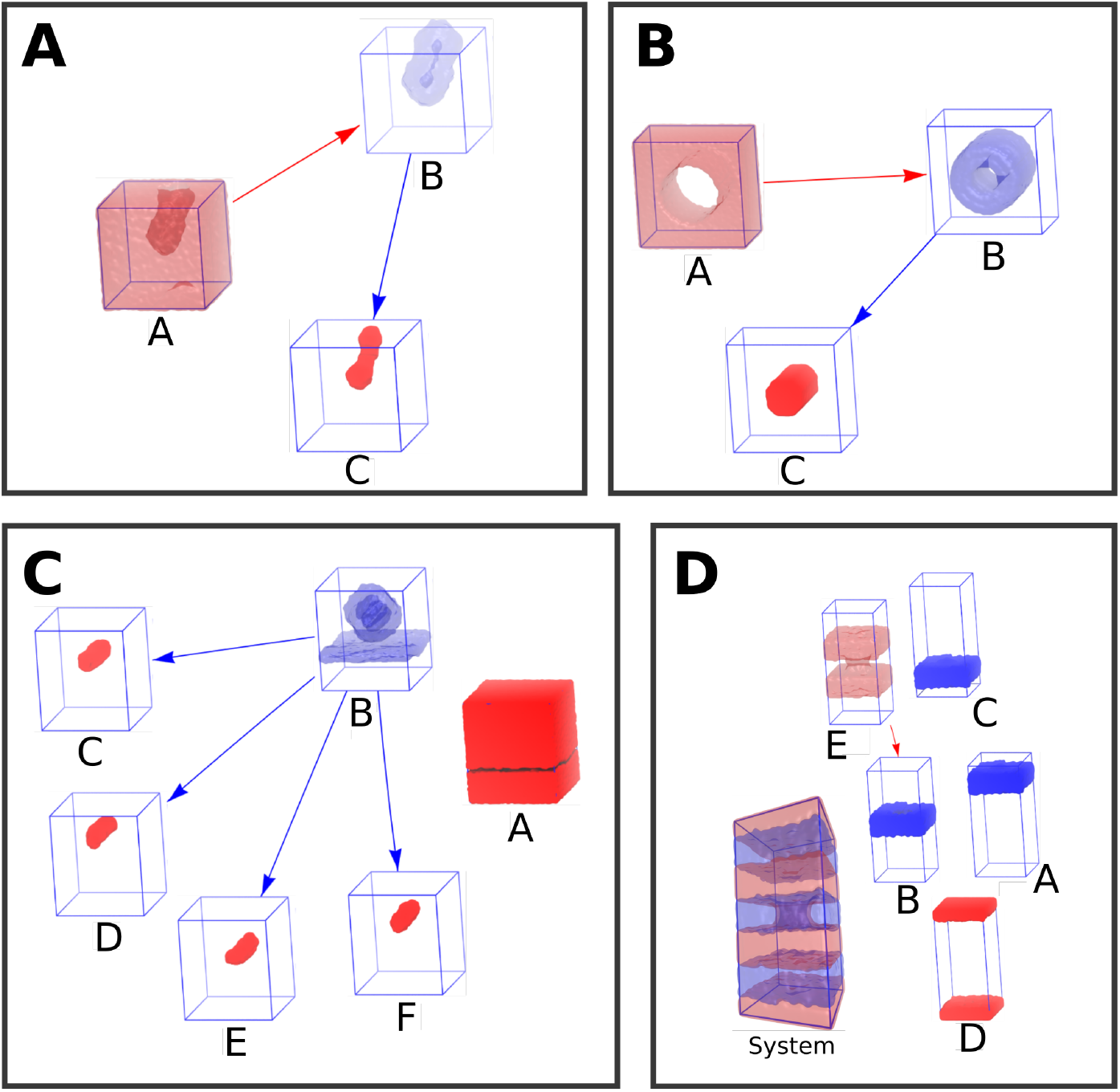
Containment hierarchy for ℝ^3^*/*ℤ^3^ MD data. (A) Acyl chains forming a vesicle. (B) Amphipathic dyes forming a periodic cylinder. (C) A lipplex fused with a bilayer, the lipoplex has four internal channels. (D) A stack of three bilayers of which one has a pore. Densities are in blue, voids are in red. Transparent densities in the containment graphs contain at least one other component.

As a first example we show the containment hierarchy for a acyl chain vesicle in solution (Fig. 11 A). There are three components in this system (_3_*A*, _0_*B*_*c*, 0_*C*_*c*_). Only component *A* is not contained, for its complement is of lower rank (_0_!*A*), the other components are contained for their complements are of higher rank (_3_!*B* and _3_!*C*). Due to their connectivity (*A*∼ *B*_*c*_, *B*_*c*_ ∼ *C*_*c*_) the resulting containment graph is linear (*A*→*B*→*C*).

The second example shows a periodic hollow cylinder in solution (Fig. 11 B). Again there are three components but this time none of them are of rank 0, meaning all components are intrinsically periodic (_3_*A*, _1_*B*_*c*, 1_*C*_*c*_). Nevertheless, only component *A* is not contained (_1_!*A*), as the others have complements of higher rank (_3_!*B* and _3_!*C*). This results in the same linear containment as is shown for the previous example (*A*→ *B*→*C*).

The third example shows a solvent phase (_2_*A*) with a lipoplex attached to a bilayer (_2_*B*). In the lipoplex there are four channels (_0_*C*_*c*, 0_*D*_*c*, 0_*E*_*c*, 0_*F*_*c*_; Fig. 11 C). Both components *A* and *B* are not contained for their ranks are equal to the ranks of their complements (_2_!*A*, _2_!*B*). All channels (*C*-*F*) are contained for they are of rank 0 (meaning their complements have to be of higher rank). Given the contacts (*A*∼ *B, B*∼ *C*_*c*_, *B* ∼*D*_*c*_, *B*∼ *E*_*c*_, *B*∼*F*_*c*_) we can assign containment, resulting in a containment graph which is not fully connected and non-linear (*A, B* →*C, B*→ *D, B*→ *E, B*→ *F*).

Finally, our last example shows a stack of three bilayers (_2_*A*, _2_*B*_*c*_ and _2_*C*), of which the central bilayer (*B*) has a toroidal pore. In between the bilayers there are two solvent phases (_2_*D* and _2_*E*), where *E* is on both sides of *B* due to the pore. All bilayer components are of rank 2, but only the complement of *B* is of higher rank (_3_!*B*), meaning *B* is contained. Given the contacts (*A* ∼ *D, A* ∼ *E, B*_*c*_ ∼*E, C* ∼ *D, C* ∼ *E*) the resulting containment graph is disconnected (*A, C, D, E* → *B*).

## 4 Conclusion

We succeeded in defining a robust and unambiguous algorithm to annotate containment in (periodic) spaces. Instead of using a winding number based approach, we use a recursive definition, making containment a non-local property of the system. The final containment is represented in a DAG where each container is capable of embedding its contained in a topological sense.

An example is the embedding of a torus in 3D periodic space. Using winding numbers we would conclude that the torus and its surrounding space are winding around each other, therefore no containment would be assigned. However, since the torus can be embedded in the surrounding space, and the surrounding space cannot be embedded in the torus, we state that the torus is contained by the surrounding space.

This was implemented in MDVContainment, an open and freely available tool which focuses on processing MD data. Although the algorithm and software can be used for non-periodic spaces, its implementation focused on the periodic cases, for this was to our knowledge still an open problem for processing MD data.

The algorithm has been demonstrated to work on both synthetic and real MD data and is actively being used for initial state building and analysis (Bentopy[8], LNPs[9], mitochondria[10], lipid morphologies[11, 12] and whole cell systems[3]), to fully automate the handling of containers. Our tool handles large systems (>100M beads) well and is mainly memory bound, where the execution times are usually less than reading the frame from drive into an MDAnalysis universe. The memory usage is related to the explicit discretization of the space and is thus related to the amount of voxels.

For any open questions or discussions we would like to point readers/users to the MDVContainment’s GitHub page (https://github.com/BartBruininks/mdvcontainment).

## 5 Acknowledgments

BMHB thanks Nastasha (the great) for the insightful bar-talks, which helped a lot in wrapping my head around the topic. BMHB also thanks Alex H de Vries, Tsjerk A. Wassenaar, Wim Couwenberg, and Erik GJ Elfving for their listening ear, and critical thinking.

## Declarations

All was said

